# A novel uncultured marine cyanophage lineage with lysogenic potential linked to a putative marine *Synechococcus* ‘relic’ prophage

**DOI:** 10.1101/325100

**Authors:** José Flores-Uribe, Alon Philosof, Itai Sharon, Svetlana Fridman, Shirley Larom, Oded Béjà

## Abstract

Marine cyanobacteria are important contributors to primary production in the ocean and their viruses (cyanophages) affect the ocean microbial communities. Despite reports of lysogeny in marine cyanobacteria, the genome sequence of such temperate cyanophages remains unknown although genomic analysis indicate potential for lysogeny in certain marine cyanophages. Using assemblies from Red Sea and *Tara* Oceans metagenomes, we recovered genomes of a novel uncultured marine cyanophage lineage, which contain, in addition to common cyanophage genes, a phycobilisome degradation protein NblA, an integrase and a split DNA polymerase. The DNA polymerase forms a monophyletic clade with a DNA polymerase from a genomic island in Synechococcus WH8016. The island contains a relic prophage that does not resemble any previously reported cyanophage but shares several genes with the newly identified cyanophages reported here. Metagenomic recruitment indicates that the novel cyanophages are widespread, albeit at low abundance. Here we describe a novel potentially lysogenic cyanophage family, their abundance and distribution in the marine environment.

**Originality-Significance Statement:** Marine cyanobacteria are major contributors to primary production in the ocean. Despite reports of lysogeny in marine cyanobacteria, genomes from lysogenic marine cyanophages have not been reported yet. Using metagenomics assemblies, we recovered complete genomes of a novel uncultured marine cyanophage lineage. Remarkably, the DNA polymerase of these uncultured phages forms a monophyletic clade with the DNA polymerase from a genomic island in *Synechococcus* WH8016. The genomic island contains a putative relic prophage that does not resemble any known cultured cyanophage but shares several genes with the newly identified cyanophage family. These findings provide both phylogenomic and abundance estimates that are missing from current ecological models of this important group of marine viruses.

## Introduction

Viruses are the most abundant biological entities on the planet, represent major genetic reservoirs and rewire their hosts metabolism (Breitbart, 2012; Paez-Espino et al., 2016). Cyanophages, viruses infecting cyanobacteria, regulate cyanobacterial communities and influence global nutrient cycles (Puxty et al., 2018).

In a previous screen of Bacterial Artificial Chromosome (BAC) libraries for photosystem II (PSII) genes, some BACs were identified as originating from cyanobacteria and others resembled known cyanophage genomes (Zeidner et al., 2005). One clone from a Mediterranean Sea sample (BAC21E04) was notable for containing numerous open reading frames (ORFs) with weak or no similarity to sequences reported in Genbank, including both non-redundant and environmental-non-redundant databases, precluding the possibility to assign affiliation to this BAC at the time. Nevertheless, BAC21E04 contains a full length viral-like D1 gene, a partial-length viral-like *talC* transaldolase gene and a putative ribonucleotide reductase (RNR) class II gene (Zeidner et al., 2005).

Viral infections can be either lytic, resulting in host lysis and virions release, or lysogenic, during which viral genomes integrate into the host chromosomes as prophages and replicate without virion production (Howard-Varona et al., 2017). Lysogeny is widespread amongst marine bacterial isolates (Stopar et al., 2003) and has been shown to occur in natural populations of marine *Synechococcus*.(McDaniel et al., 2002). Among the benefits and consequences of lysogeny, reviewed elsewhere e.g. (Howard-Varona et al., 2017), it has been proposed to play a role in the protection phages from decay (Breitbart, 2018), contribution to host survival under unfavorable environments by suppression of unneeded metabolic activities (Paul, 2008), and phage-mediated horizontal gene transfer of bacterial DNA (Chen et al., 2018). Although temperate phages infecting either freshwater cyanobacteria *Anacystis nidulans* (Lee et al., 2006) or marine filamentous cyanobacteria (Ohki and Fujita, 1996) are known, to date prophage induction in marine cyanobacteria has only been studied using natural populations or the cultured *Synechococcus* GM 9914 (McDaniel and Paul, 2005; McDaniel et al., 2006). *Synechococcus* GM 9914, isolated from the Gulf of Mexico, in a non-axenic culture was shown to produce virus-like particles (VLP) containing single stranded DNA under exposure to mitomycin C or high-light stress, 300% more light than the normal culture level, unfortunately the DNA of these VLPs was not sequenced (McDaniel et al., 2006).

Here, we report the identification of a novel, widespread lineage of uncultured cyanophages related to BAC21E04. These cyanophages share properties related to temperate phages and shares synteny with a putative relic prophage in *Synechococcus* WH8016, suggesting that these uncultured cyanophages are potentially lysogenic.

## Results and Discussion

To increase our knowledge regarding uncultured cyanophages, we examined metagenomic assemblies from the Red Sea (Philosof et al., 2017) and *Tara* Oceans expedition microbiomes and viromes (Brum et al., 2015; Sunagawa et al., 2015). Four contigs (Supporting Information File 1) of putative viral genomes related to BAC21E04 were identified. These contigs, of about 86 kbp (Supporting Information File 2), contain overlapping terminal regions suggesting that they represent complete, terminally redundant viral Metagenome Assembled Genomes (MAGs). The four MAGs contained a cyanophage-like ribonucleotide reductase (RNR) class II gene (Supporting Information Figure 1 and Supporting Information File 3), a *talC* transaldolase gene (Supporting Information Figure 1 and Supporting Information File 4), and an exonuclease (*exo*). Three MAGs carry tRNA genes, an integrase encoding gene, and a *psbA* gene with viral signatures (Sharon et al., 2007) (Supporting Information File 5). Also, detected for the first time in marine cyanophages, genes coding for the phycobilisome degradation protein NblA were found in each of the newly identified cyanophages, and three of the MAGs contain two different copies of *nblA*. Single *nblA* genes have been previously reported in freshwater cyanophages (Gao et al., 2012; Ou et al., 2015) that infect cyanobacteria using phycobilisomes for light harvesting. Phycobilisomes, large extrinsic multisubunit light-harvesting complexes typical of most cyanobacteria, have been found in *Synechococcus* but not in *Prochlorococcus* (Dufresne et al., 2003). However, as functional, infection-expressed phycobilisome pigments biosynthesis genes have been characterized in *Prochlorococcus* cyanophages (Dammeyer et al., 2008) we cannot assume that the putative host of these *nblA* carrying cyanophages is *Synechococcus*. The terminase large subunit (TerL), portal, major capsid protein (MCP), and putative tail proteins were identified based on structural predictions. However, no other known phage structural proteins could be detected as protein identity was too low for alignment-based methods to work.

All four MAGs were analyzed using VirFam (Lopes et al., 2014), which classifies phages into *Myoviridae, Podoviridae* or *Siphoviridae* according to their neck protein organization. VirFam identified the predicted MCP, Portal and TerL ORFs, however it could not classify them into any of its neck-type categories, suggesting that these cyanophages could belong to a new lineage (Supporting Information File 6).

Due to the lack of good alignments for the traditionally used “phage marker genes” such as portal, MCP or terL protein we employed a strategy similar to the one used by (Rohwer and Edwards, 2002; Mizuno et al., 2013) for the classification of phages based on total shared core pangenome. The gene content of the MAGs and 82 genomes from cyanophages infecting *Synechococcus* and *Prochlorococcus* (Supporting Information File 7) was analyzed identifying 14,564 proteins that cluster into 2,376 orthologs. Analysis of the shared orthologs between viruses allowed to delineate four clusters of cyanophages corresponding to the viral families *Myoviridae, Podoviridae, Siphoviridae* and one composed exclusively of the MAGs from this study (Fig. 1a). A similar pattern was achieved when genomic nucleotide distances were used for clustering (Supporting Information Figure 2). The clustering results suggested that the MAGs presented here belong to a new cyanophage lineage whose closest relatives are the unclassified freshwater *Synechococcus* cyanophage S-EIV1 (50% shared core proteome) and the putative siphovirus S-SKS1 (20% shared core proteome). In our core proteome analysis, S-SKS1 clusters among the *Myoviridae* group, such re-classification has also been reported before (Roux et al., 2016).

**Figure 1.**
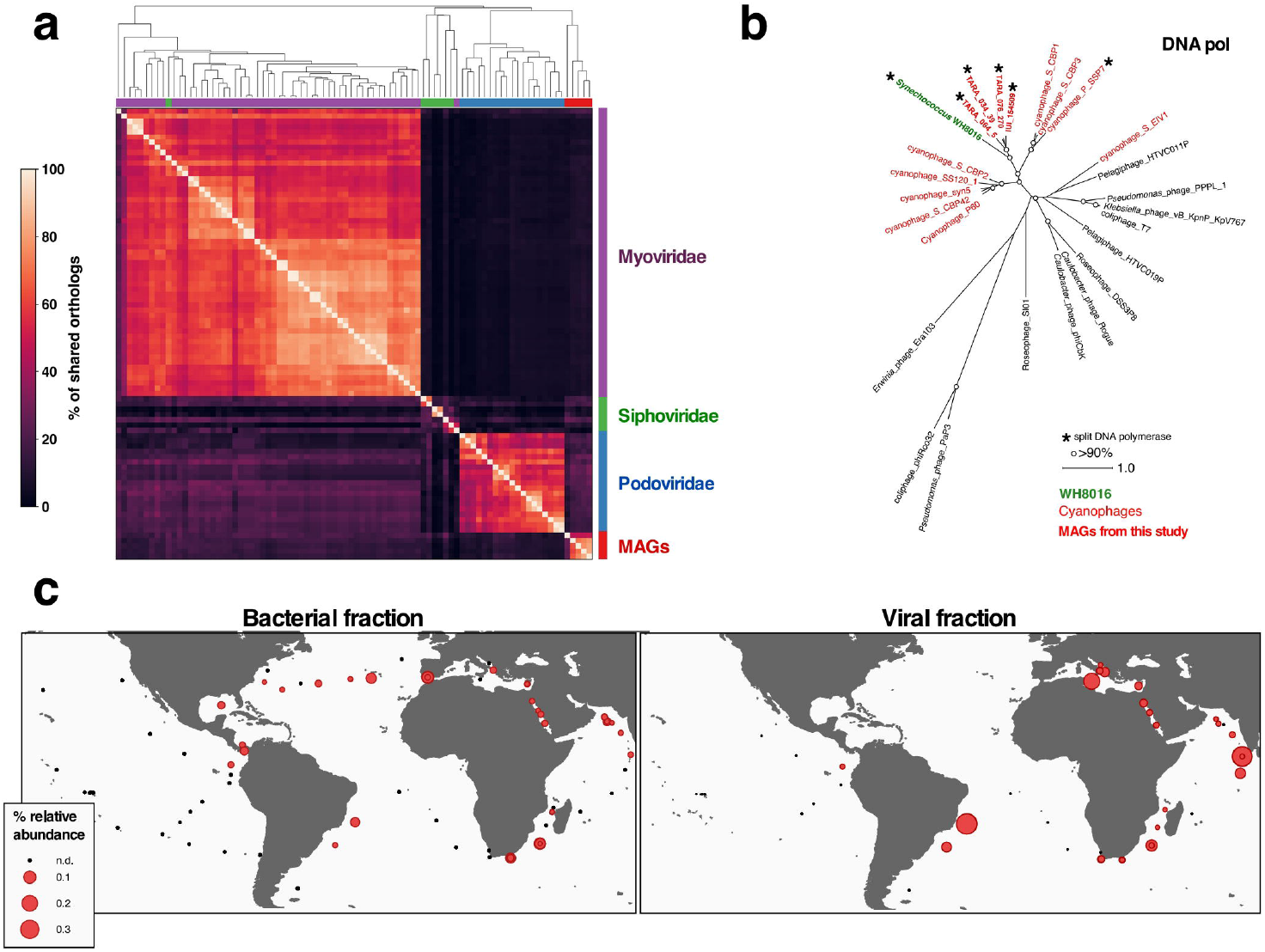
**a)** Comparison of Cyanophages genomes clustered according to percentage of shared orthologs. Heatmap representation of protein orthologs shared by cyanophages, scaled between 0 (darker) and 100% (lighter). Dendrogram tips are colored according to the taxonomic affiliation of the cyanophages using the following key: *Myoviridae* (purple), *Podoviridae* (blue), *Siphoviridae* (green), and the MAGs identified in this study (red). Clustering identifies four large groups corresponding to the taxonomic families from the cyanophages (full details in the supporting information). **b)** Phylogenetic protein tree of DNA Polymerase. The newly identified MAGs and *Synechococcus* WH 8016 are shown in bold red. Cyanophage and cyanobacteria sequences are shown in red and green, respectively. Split DNA polymerases are marked with an asterisk. The MAGs form a monophyletic clade with a split DNA polymerase from a genomic island in *Synechococcus* WH8016. **c)** Distribution and relative abundance of the newly identified cyanophages in Tara Oceans metagenomes. Each red circle is scaled to represent the relative abundance of the MAGs in each station. Black points indicate stations where the MAGs could not be detected (<0.01%).

Metagenomic recruitment of the *Tara* Oceans metagenomic reads into the MAGs was used to determine their coverage (Supporting Information Figure 3), presence-absence of their genes (Supporting Information Figure 4) as well as abundance estimation in the *Tara Oceans* datasets (Supporting Information File 8). The mapping results indicated the presence (>0.01% relative abundance) of these cyanophages in 36 samples from 16 stations with an average of 0.07% (Fig. 1c). A viral fraction (<0.2 μm) sample from the South Atlantic (TARA_076) had the highest abundance of the new cyanophages (0.39%), suggesting the presence of cell-free virions from the new family in the environment. To assess if the viral fraction samples where the MAGs were recruited had cyanobacterial contamination, we estimated cyanobacterial abundance in those samples. Cyanobacterial abundance was estimated using pseudo-mapping with Salmon (v0.12.0-alpha) and a collection of cytochrome b6 (*petB*) sequences (Supporting Information File 9). *petB* has been used as a marker gene for marine cyanobacteria metagenomic abundance estimation (Farrant et al., 2016). Salmon results showed no detection of *petB* on any of the virome samples, indicating that cyanobacterial contamination in these samples is improbable. However, 24 of the 36 samples where the MAGs were detected correspond to the bacterial fraction (0.2-3.0 μm) probably indicating ongoing infections during sampling (Philosof et al., 2017).

The new cyanophages also encode a split DNA polymerase, which forms a new cluster within the T7-like cyanophage DNA polymerase family (Fig. 1b). Unexpectedly, a split DNA polymerase from *Synechococcus* WH8016 also clustered with the new phage family (Supporting Information File 10). Examination of the vicinity of the DNA polymerase genes in the genome of WH8016 revealed the existence of a previously uncharacterized genomic island of approximately 70 kbp (Fig. 2). The genomic island is delimited by a tRNA-Pro and a tRNA-Arg and contains several genes (such as split DNA polymerase genes, integrase, phage/plasmid related protein; Fig. 2) in synteny that exhibit similarity with ORFs from the new viral lineage (Fig. 3).

**Figure 2.**
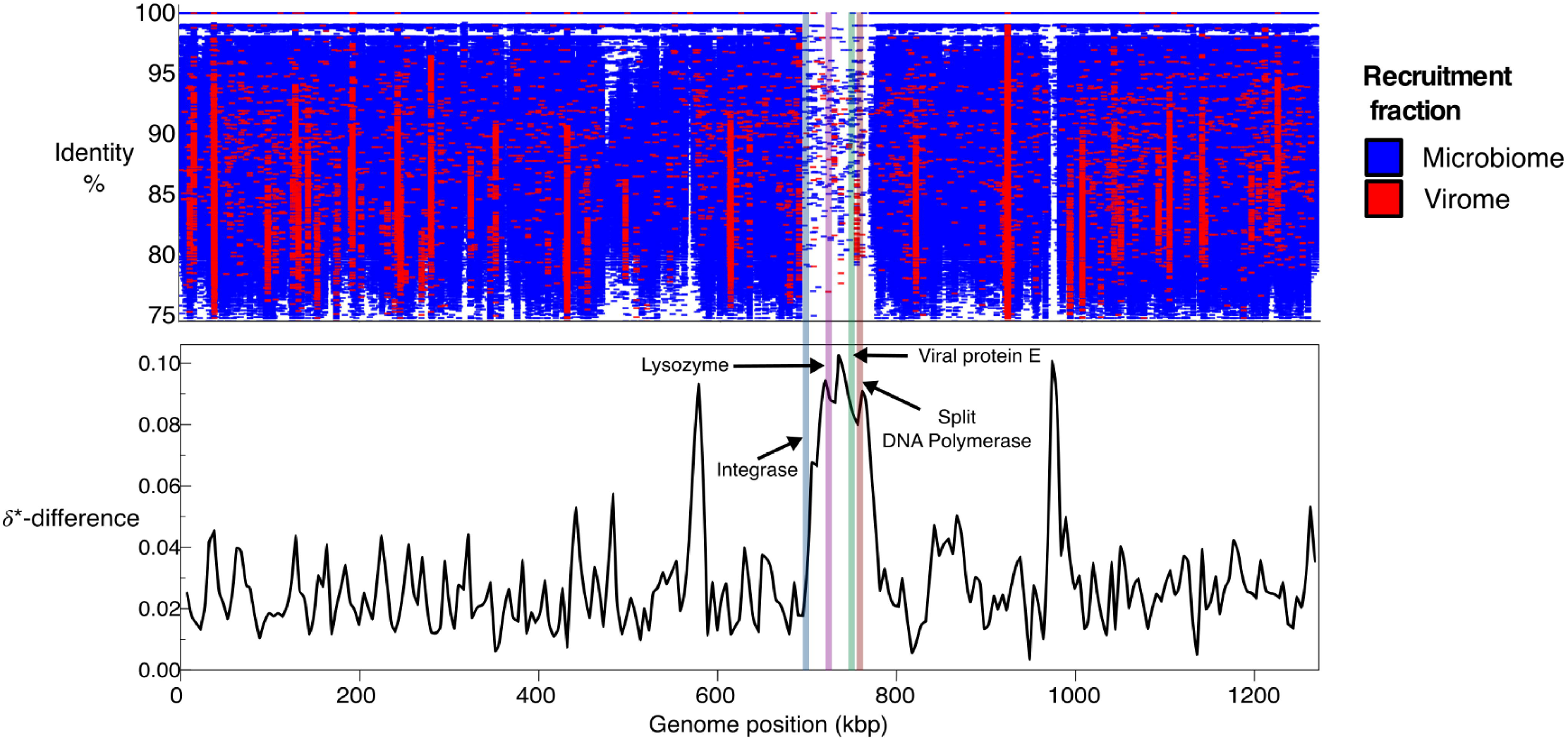
A genomic island in the genome of *Synechococcus* WH8016. Top: Recruitment of *Tara* Oceans reads from 331 metagenomes onto *Synechococcus* WH8016 genome. Each recruited read is drawn according to the position in the genome where it aligned and the nucleotide identity to the recruitment area. Reads are colored according to the sample fraction of origin, red for virome (<0.2 μm) and blue for microbiome (0.2–3.0 μm). Bottom: Karlin signature (*δ*\*) difference, the difference in the relative abundance of dinucleotides between a sliding window and the whole sequence (Karlin et al., 1998), was calculated using a window size of 10000 bp and a step size of 5000 bp. Colored bars spanning both panels indicate position of some of the shared proteins in synteny with the newly identified cyanophages.

**Figure 3.**
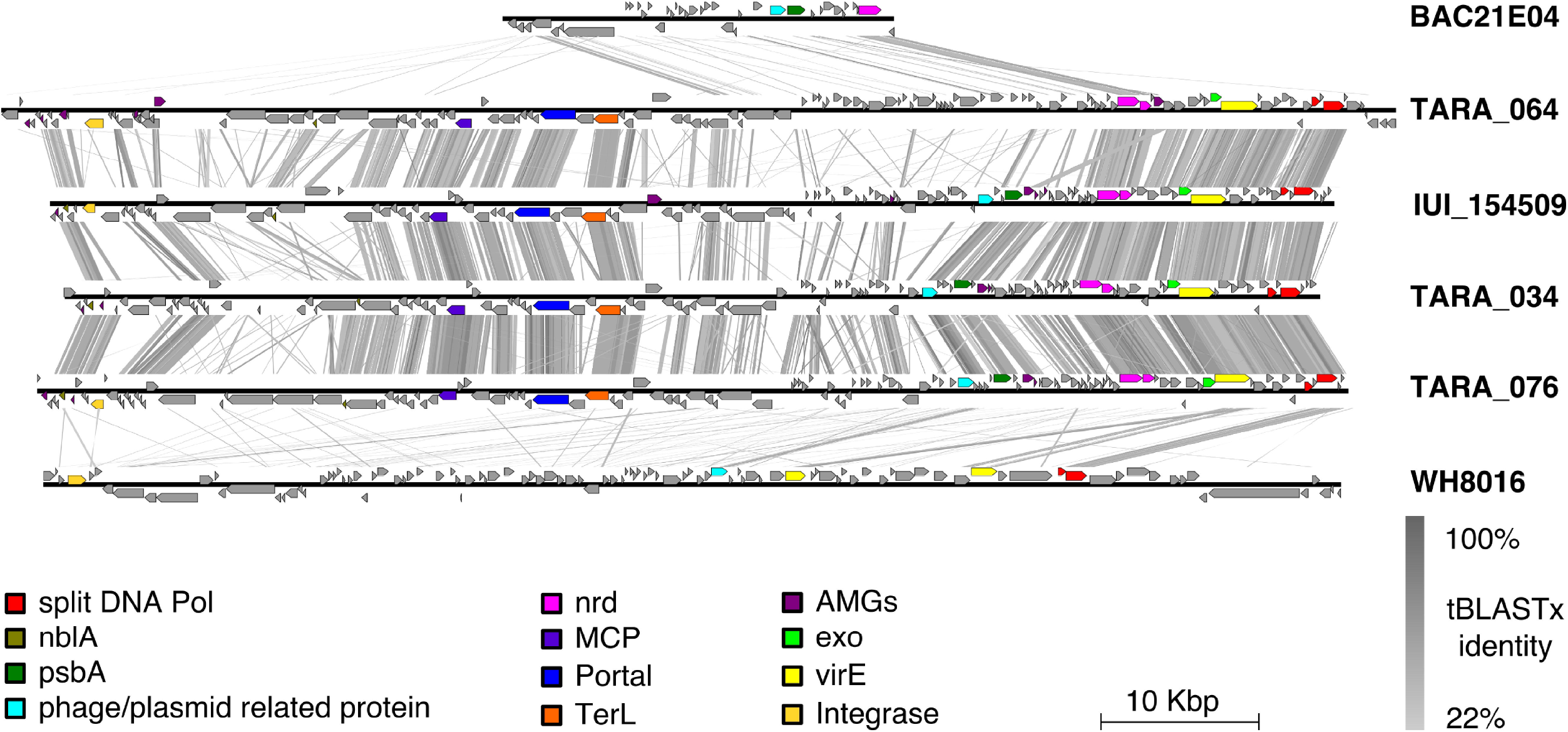
Synteny plots for the MAGs from the reported viral lineage. Distinctive viral genes, cyanophage marker genes and genes of interest are colored according to the legend. Using tblastx regions of similarity among the phages, BAC21E04 and the genomic island of WH8016 were identified. The synteny plot was obtained using Easyfig (Sullivan et al., 2011) and the origin of the MAGs was readjusted for clarity. The figure was completed using the open-source vector graphics editor Inkscape (available from http://inkscape.org/).

Neither production of viral particles nor expression of genes inside the genomic island could be detected, with TEM and qPCR respectively, after exposure of WH8016 to mitomycin C at 5 μg/mL. Although the concentration of mitomycin C employed in the assay was five times higher than previous reported ones for the marine environment (1 μg/mL) (Paul and Weinbauer, 2010; Knowles et al., 2017). Moreover, we also tried induction of WH8016 using heat-shock, darkness exposure, or co-cultivation with other marine cyanobacteria (see Methods) without detection of lysis plaques.

Prophages can also be induced by exposure to different environmental stimuli as pollutants, metals, sunlight, UV irradiation, nutrient deficiency, or pressure changes (Jiang and Paul, 1994; Cochran et al., 1998; McDaniel et al., 2002; Aertsen et al., 2004; Lee et al., 2006). Additionally, induction of some lysogenic freshwater cyanophages in laboratory conditions requires the use of mitomycin C at 20 μg/mL (Dillon and Parry, 2008) and therefore the mitomycin C concentration and the environmental stressors used in this study might not have been enough to produce a SOS response in WH8016.

Prophages can undergo “domestication” processes driven by their hosts looking to select for beneficial viral traits while losing genes like those encoding structural and lysis components (Bobay et al., 2014; Howard-Varona et al., 2017). These can eventually result in relic prophages and genomic islands that facilitate chromosomal insertions while providing resistance to other phages (Schwartz and Lindell, 2017) and could influence the expression of neighboring genes (Howard-Varona et al., 2017). Furthermore, the variety of bacterial mobile genetic elements include several examples of entities which can only be mobilized by a helper phage. These entities include phage-inducible chromosomal islands (PICIs) and phage-inducible chromosomal island-like elements (PLEs) (Fillol-Salom et al., 2018). PICIs and PLEs have sizes of up to 19kb, do not possess identifiable regulatory, replication or packaging modules and require the help of an additional phage to excise from the bacterial chromosome (Fillol-Salom et al., 2018). Therefore, under the conditions of our assays and the lack of recognizable viral structural proteins, we conclude that the genomic island in *Synechococcus* WH8016 represents either a non-mitomycin C inducible phage, an element which requires the help of another phage to mobilize or a putative ‘relic’ prophage that lost the ability to enter into lytic cycle.

Based on our observations, we suggest that the newly identified marine phages form a new, potentially lysogenic, cyanophage family related to the newly characterized ‘relic’ prophage in *Synechococcus* WH8016. The identification of putative lysogenic SAR11 phages (Zhao et al., 2018) and other recent studies suggest that lysogeny in the marine environment could be implicated in ecological processes on broad scales (Knowles et al., 2016; Wigington et al., 2016; Knowles et al., 2017).

Although almost half of the sequenced bacteria are lysogens (Touchon et al., 2016), available marine cyanobacterial genomes do not contain complete prophages in them. The lack of recognizable prophages in marine cyanobacterial genomes could be a result of multiple factors among which we can enlist two. First, the relatively low number of cyanobacterial genomes available in public databases when compared to other bacterial phyla (Alvarenga et al., 2017) which makes difficult the use of automated prophage detection tools. Second, in accordance to what has been proposed for the recently described temperate pelagiphages (Zhao et al., 2018), many marine cyanobacteria possess a small cell size and streamlined genome architecture (Larsson et al., 2011) which could impose an additional cost to carry prophages (Zhao et al., 2018) and is associated with a low proportion of prophages (Touchon, 2016). Genomic evidence for a partial prophage in *Prochlorococcus* cells (Malmstrom et al., 2013) and a relic prophage identified in marine *Synechococcus* RS9917 (Sullivan et al., 2009), are examples of unstable integrations of prophages in marine cyanobacteria.

Along with the abundance estimates for the novel lineage of cyanophages reported here, we suggest that lysogeny in cyanobacteria might be more widespread than previously thought, adding to the range of life styles that marine cyanophages exhibit.

## Methods

### Re-assembly of *Tara* Oceans data

Raw reads for the 244 *Tara* Ocean microbiome (Sunagawa et al., 2015) and viromes (Brum et al., 2015) samples were downloaded and assembled separately using the idba-ud (Peng et al., 2012) and megahit v1.0.4 (Li et al., 2015) assemblers. idba-ud was used with default parameters, for megahit the ‘--presets meta-sensitive’ option was used. Only scaffolds >=500 bp were considered. To correct mis-assemblies we used a read-mapping based in-house pipeline that consists of two steps. First, the pipeline used read mapping information to identify short mis-assemblies such as wrong base calls and short missing segments. Next, it used paired read mapping to identify scaffolding errors (wrongly connected contigs) and to fill gaps (stretches on Ns). To fill gaps the pipeline used a local assembly approach in which reads that map to the area of the gap and they paired-end reads are assembled and adjusted to the region. For read mapping we used bowtie2 (Langmead et al., 2012) with the ‘— sensitive’ option.

### Identification of contigs related to BAC21E04

Open reading frames (ORFs) from BAC21E04 (Zeidner et al., 2005) were used as query for tBLASTx v2.6.0 (Camacho et al., 2009), with default parameters, against the re-assembled contigs from *Tara* Oceans and the Red Sea assemblies (Philosof et al., 2017)(NCBI BioProjectID: 362713: Red Sea Diel), four contigs (Supporting Information File 1) with sizes above 79 kbp were identified and selected. These contained overlapping terminal regions suggesting that they represent complete, terminally redundant viral Metagenome Assembled Genomes (MAGs). Blastn was used to detect overlapping terminal repeats. The first and last 1000 bp of each one of the MAGs were used as query for a blastn analysis with default parameters against the re-assembled Tara Oceans metagenomes. Hits with >=99% identity were aligned to the blastn queries using Sequencher v5.0.1 and UGENE.

### MAGs annotation

ORFs in the MAGs were identified using prodigal v2.6.3 (Hyatt et al., 2010). Annotation of the ORFs was performed first using prokka v1.12 (Seemann, 2014). The annotation with prokka is based on blastp (Camacho et al., 2009) and hmmer (Eddy, 2011) using the default prokka databases for Viruses and Prokaryotes. Prokka assigned putative function in average to 10% of the ORFs for each MAG. ORFs without annotation or annotated as hypothetical proteins were analyzed using hmmscan v3.1b2 (Eddy, 2011) with the pVOG database (Grazziotin et al., 2017) (cutoff for e-value 1e-15), interproscan (Quevillon et al., 2005), pfam_scan.pl v1.6 (using --cut_ga and a HMM coverage cutoff value of 0.35) against Pfam31 (Finn et al., 2006) and the hhpred pipeline from the hhblits suite (Remmert et al., 2011) (cutoff for probability: 85%).

### Orthologs detection and clustering

To estimate the relationship in gene content between the MAGs and a collection of 82 genomes of cyanophages infecting *Prochlorococcus* or *Synechococcus*, OrthoFinder v2.2.0 (Emms and Kelly, 2015) was used with the option ‘-S blast’. All 14564 proteins from 86 genomes (4 MAGs + 82 cyanophages) were clustered into 2376 orthologs. The number of orthologs shared by each pair of cyanophages, contained in the file Orthogroups_SpeciesOverlaps.csv produced by OrthoFinder, was normalized per row. The normalized results were then used to generate a heat map, clustered using clustermap from the python package seaborn v0.8.1 (available at: https://zenodo.org/record/883859).

### Genomic distance estimation

Comparison of the MAGs at nucleotide level against the collection of cyanophage genomes previously mentioned was performed using mash (Ondov et al., 2016) with a sketch size of 7500. Genomic distances obtained by mash were used as input for clustermap from seaborn v0.8.1.

### MAGs abundance estimation

MAGs were used for metagenomic recruitment of the Tara Oceans microbiome and viromes (n=399) using Bowtie2 v2.3.0 (Langmead and Salzberg, 2012). A Bowtie2 index was built using a concatenated FASTA file containing the sequences of the 4 MAGs and the 82 cyanophages. The index was used to map the short reads from the datasets (bowtie2 parameters: --no-unal) and the results were stored as indexed BAM files using samtools v1.3.1(Li et al., 2009). Anvi’o v4.0 (Eren et al., 2015) metagenomic workflow (Delmont and Eren, 2018) was followed to estimate the MAGs relative abundance from the metagenomic samples. Briefly, the FASTA file containing the cyanophage genome sequences used to build the Bowtie2 index was used to generate an Anvi’o contigs database, contigs.db, using the program ‘anvi-gen-contigs-database’ with default parameters. Each indexed BAM file was processed using the program ‘anvi-profile’ and the contigs.db file generated in the previous step to generate an anvi’o profile database. The anvi’o profiles were then merged using anvi-merge with the parameter ‘-- skip-concoct-binning’. Using ‘anvi-import-collection’ each of the contigs in the merged profile were linked to the genome they originated from. Finally, the contigs belonging to the MAGs in the merged anvi’o profile were summarized using ‘anvi-summarize’. The file relative_abundance.txt from the anvi’o summary was used to generate the distribution map as specified in the visualization section below. The Anvi’o summary for the new cyanophages in Tara Oceans samples, containing the abundance estimations, can be accessed at: https://osf.io/q3khc/

### Cyanobacteria abundance estimation

Relative abundance of cyanobacteria was calculated using Salmon (v0.12.0-alpha) (Patro et al., 2017). A collection of 649 cytochrome b6 (*petB*) DNA sequences (Supporting Information File 9) from photosynthetic microorganisms (chloroplasts, freshwater and marine cyanobacteria) was used to create a Salmon index. Occurrence of petB in the 399 metagenomes from the Tara Oceans microbial, giant viruses and viral fractions was quantified with the index using Salmon in the quasi-mapping mode with the following parameters ‘--meta --incompatPrior 0.0 -- seqBias –gcBias --numAuxModelSamples 2500000 --numBootstraps 100 -- validateMappings’. Quantification results were processed by tximport (v1.10.0) (Soneson et al., 2015), followed by normalization with edgeR (v3.24.2) (Robinson et al., 2010). Reads per kilobase per million were calculated from the normalization results by the edgeR function ‘edgeR::rpkm’. Abundance results were processed in Python (v3.7) and pandas (version 0.23.4) (McKinney, 2010). The Salmon index as well as the quantification results for each sample can be found at: https://osf.io/q3khc/

### Metagenomic recruitment using Synechococcus WH8016

*Tara* Oceans microbiome and virome metagenomic datasets were mapped against the sequence of *Synechococcus* WH8016 (GenBank: AGIK01000001) using Bowtie2. BAM files containing the mapping results were parsed to extract the mapped reads alignment position and nucleotide identity. Only reads with at least 75% identity were retained. The alignments data was used to generate the recruitment plot.

### NblA screening of cyanobacterial genomes

All the coding DNA sequences from the order *Synechococcales* were downloaded from RefSeq (Retrieved on 2017-03-29) and used for query in a hmmscan against a HMM database composed by the NblA pfam model (PF04485; available at http://pfam.xfam.org) and the NblA predicted ORFs from the MAGs. No hit to the NblA HMM could be identified in sequences coming from marine cyanobacteria. The NblA hmmer database can be downloaded at: https://osf.io/q3khc/

### Karlin signature difference of *Synechococcus* WH8016

The genome of WH8016 (GenBank accession: AGIK01000001.1) was analyzed by calculating the Karlin signature (*δ*\*) difference, the difference of the relative abundance of dinucleotides between a sliding window and the whole sequence (Karlin et al., 1998), using with UGENE (Okonechnikov et al., 2012) with a window size of 10 000 bp, a step size of 5 000 bp and exported as bitmap.

### Phylogenetic trees construction and analysis

Amino-acid sequences of the genes coding for DNAPol, RNR and TalC in the MAGs were aligned to related sequences from picocyanobacteria, phages and cyanophages retrieved from GenBank. The alignment and phylogenetic analysis of each set of protein sequences was performed using the http://www.phylogeny.fr/ pipeline (Dereeper et al., 2008) with default parameters according to the following workflow: First, amino-acid sequences were aligned using MUSCLE (Edgar, 2004), then the alignment was curated using Gblocks (Castresana, 2000). Finally phylogenetic trees were inferred by PhyML (Guindon et al., 2009; Guindon et al., 2010) using 100 bootstraps. The sequences used for the alignment can be retrieved at: https://osf.io/q3khc/

### Visualization

The map (Fig. 1c) was generated using the library ggplot2 (Wickham, 2009) for R. The clustermap and lmplot functions from the python package seaborn v0.8.1 (available at: https://zenodo.org/record/883859) were used to draw the Heat maps (Fig. 1a) and recruitment plot (Fig. 2) respectively. All figures were formatted for publication using the open-source vector graphics editor, Inkscape (available at: http://inkscape.org/)

### Data and code availability

The sequence of the MAGs can be found in Supporting Information File 1. The rest of the sequences are deposited at GenBank as follows: BAC21E04 [AY713439-AY713444], *Synechoccocus* WH8016 [AGIK01000001]. Supporting Information File 7 contains the accessions for the collection of cyanophages. Supporting data can be retrieved from https://osf.io/q3khc/ while the Jupyter Notebooks and scripts used are publicly available at https://github.com/BejaLab/TM_Cyanophages.

### Strains and culture conditions

*Synechococcus* spp. strains WH8016, WH7803, WH7805, WH8102, WH8109, and WH5701 were grown in artificial seawater-based medium (ASW) (Wyman et al., 1985) modified as described previously (Lindell et al., 1998). Cultures were grown at 21 °C under cool white light with a 14:10 h light:dark cycle at an intensity of 10–14 μmol photons m-2 s-1 during the light period, unless otherwise stated.

### Mitomycin-C induction

Experiments with Mitomycin-C were performed according to (Zhao et al., 2010) with some modifications. At mid log phase a culture of *Synechococcus* WH8016 was treated with 5 μg mL^−1^ of mitomycin C. After 48h cells were pelleted and the supernatant was concentrated using a 0.22μm PVDF filter unit. Samples for transmission electron microscopy (TEM) were prepared according to (Sabehi et al., 2012).

### Quantification of DNA by real time qPCR

*Synechococcus* WH8016 cultures at mid log phase were treated with 5 μg mL^−1^ of mitomycin C. Samples were collected at 0, 2, 4, 8, 12 and 24 h after exposure to mitomycin C. A 9 mL volume of treated or untreated cells was filtered onto 0.22 μm Durapore filters (Millipore). The filters were transferred to tubes containing 1 mL RNAlater (Ambion), frozen, and stored at −80 °C. Total RNA was extracted using a mirVana RNA isolation kit (Ambion). Genomic DNA was removed using a Turbo DNA-free kit (Ambion). cDNA was made using a High-Capacity cDNA Reverse Transcription Kit (Applied Biosystems). Integrase and DNApol genes from WH8016 were chosen for transcript analysis by real-time PCR.

Each real-time PCR reaction contained 1x Roche universal probe library (UPL) master mix (LightCycler 480 Probes Master, Roche), 80 nM UPL hydrolysis probe, 500 nM primers and cDNA template in a total volume of 25 μL. Reactions were carried out on a LightCycler 480 Real-Time PCR System (Roche). The program used was one step of 95 °C for 15 min followed by 45 cycles of amplification, each including 10 s denaturation at 95 °C and a 30 s combined annealing and elongation step at 60 °C, at the end of which plate fluorescence was read (FAM; Ex/Em 465/510 nm). The primers used in real-time PCR assays is provided in the Supporting Information File 11.

### Cocultivation with marine cyanobacterial strains

*Synechococcus* WH8016 was cocultured in ASW plates with *Synechococcus* spp. strains WH7803, WH7805, WH8102, WH8109, and WH5701. The plates were kept for one month after a bacterial lawn was formed and examined looking for viral plaques.

### Cultivation under stress conditions

*Synechococcus* WH8016 cultures were grown on ASW plates under standard conditions and then exposed to a series of environmental stressors for two weeks. Heat stress: From 21 to 37 °C. High light stress: From 14:10h L/D and 10–14 μmol photons m^−2^ s^−1^ to continuous light and 20 μmol photons m^−2^ s^−1^. Light deprivation: After growing on L/D photoperiod the plates were covered from light. After growing for two weeks under the stress the plates were examined, looking for viral plaques.

## Author Contributions

J.F.-U. and O.B conceived the project. J.F.-U., A.P., I.S., and O.B. performed bioinformatic analyses. J.F.-U., S.F. and S.L. performed mitomycin C experiments. J.F.-U. and O.B. wrote the manuscript with contributions from all authors to data analysis, figure generation, and the final manuscript.

## Acknowledgements

We thank Laurence Garczarek for sharing unpublished data for *Synechococcus* WH8016, and Curtis Suttle and Caroline Chénard for sharing DNA polymerase sequences. This work was funded by a European Commission (ERC Advanced Grant no. 321647), and the Louis and Lyra Richmond Memorial Chair in Life Sciences (to O.B.).

## Supporting Information Figure legends

**Supporting Information Figure 1.** Phylogenetic protein trees: Transaldolase and RNR. The newly identified contigs and *Synechococcus* WH8016 are shown in bold. Cyanophage and cyanobacteria sequences are shown in red and green, respectively.

**Supporting Information Figure 2.** Clustering of genomic distance among cyanophages infecting *Synechococcus* and *Prochlorococcus* using Mash (Ondov et al., 2016). The heat maps illustrate the pairwise similarity as genomic distance between genomes, scaled between 0 (darker) and 1 (lighter). Genome groups are colored in the dendrogram according to the taxonomic group of the cyanophages using the following key: *Myoviridae* (purple), *Podoviridae* (blue), *Siphoviridae* (green), and the MAGs identified in this study (red).

**Supporting Information Figure 3.** Nucleotide level MAGs Tara Oceans coverage. The coverage of each nucleotide position from the MAGs in the Tara Oceans metagenomes was recovered from bowtie2 mappings. On top of each plot a schematic representation of the ORFs genomic maps for each MAG is presented, colored as indicated in the legend.

**Supporting Information Figure 4.** Anvi’o detection of the MAGs ORFs in the Tara Oceans metagenomes. In box-plots detection values for each ORF in samples with at least 75% mean MAG detection. Boxes colored according to the sample fraction. Mean 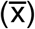 and median (M) displayed next to the corresponding sample.

## Supporting Information Legends

**Supporting Information File 1.** Sequences in FASTA format for the MAGs identified in the study.

**Supporting Information File 2.** Genometrics of the MAGs. Properties of the MAGs identified in the study: Length, %GC content, Number of Open Reading Frames, Position of Open Reading Frames of Interest

**Supporting Information File 3.** Amino acid alignments in FASTA format used for the RNR phylogenetic tree.

**Supporting Information File 4.** Amino acid alignments in FASTA format used for the TalC phylogenetic tree.

**Supporting Information File 5.** Annotation table. List of the ORFs and tRNAs identified in the MAGs with putative annotation, method of annotation, DNA sequence and amino acid sequence.

**Supporting Information File 6.** VirFam results for the classification of the MAGs.

**Supporting Information File 7.** Cyanophages used for cluster comparisons. List of 86 cyanophage genomes used in the cluster comparison analysis, contains 82 sequenced cyanophages deposited at NCBI GenBank and the 4 MAGs from the study.

**Supporting Information File 8.** Coverage, detection, and abundance summaries calculated with Anvi’o of the MAGs across the Tara Oceans metagenomes.

**Supporting Information File 9.** Accessions of the *petB* sequences used for the cyanobacterial abundance estimation.

**Supporting Information File 10.** Amino acid alignments in FASTA format used for the DNAPol phylogenetic tree.

**Supporting Information File 11.** Primers used for qPCR detection of genes inside the genomic island of WH8016.

