## Supplementary figures and images for "A novel uncultured marine cyanophage lineage with lysogenic potential linked to a putative marine *Synechococcus* ‘relic’ prophage"

### Supplemental Figure 1

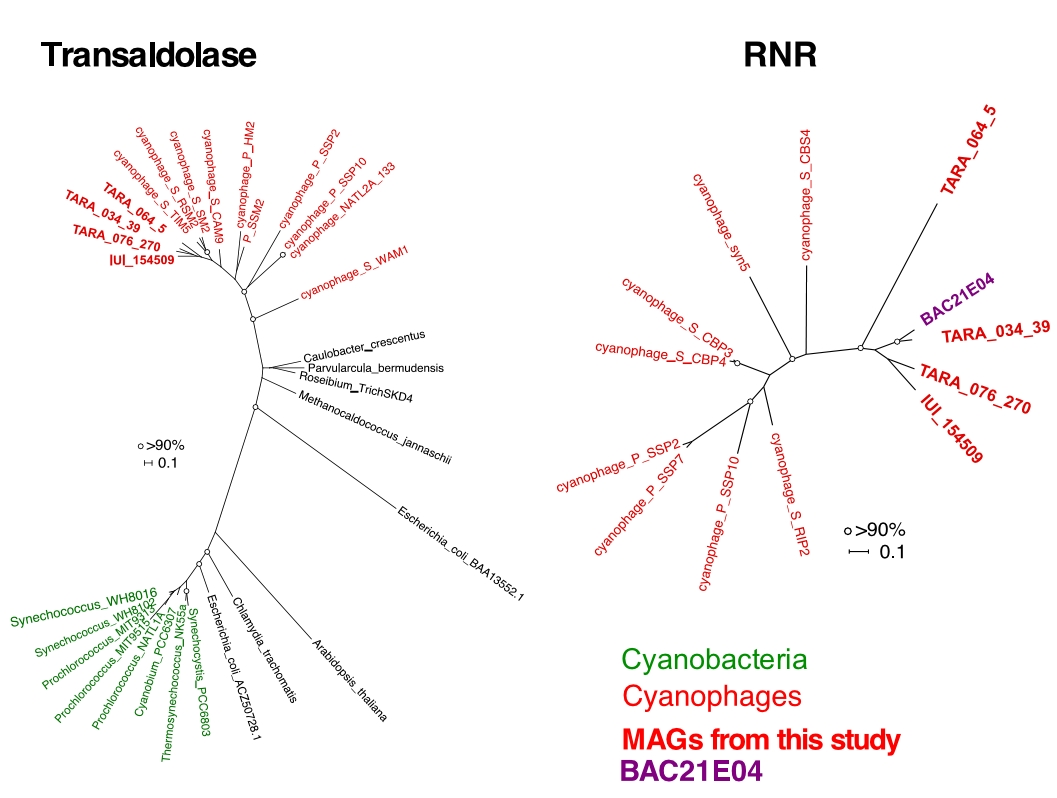

### Supplemental Figure 2

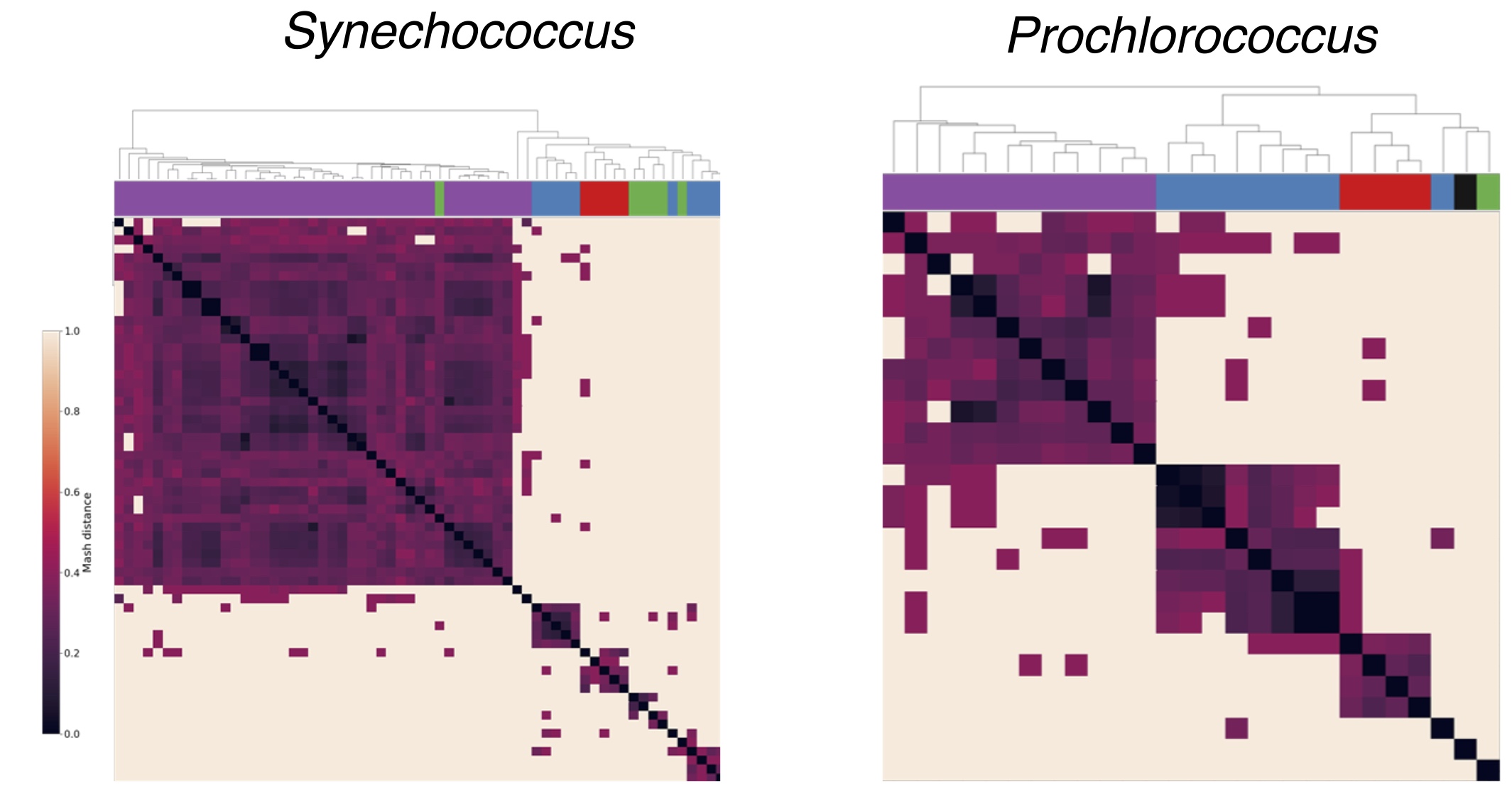

### Supplemental Figure 3

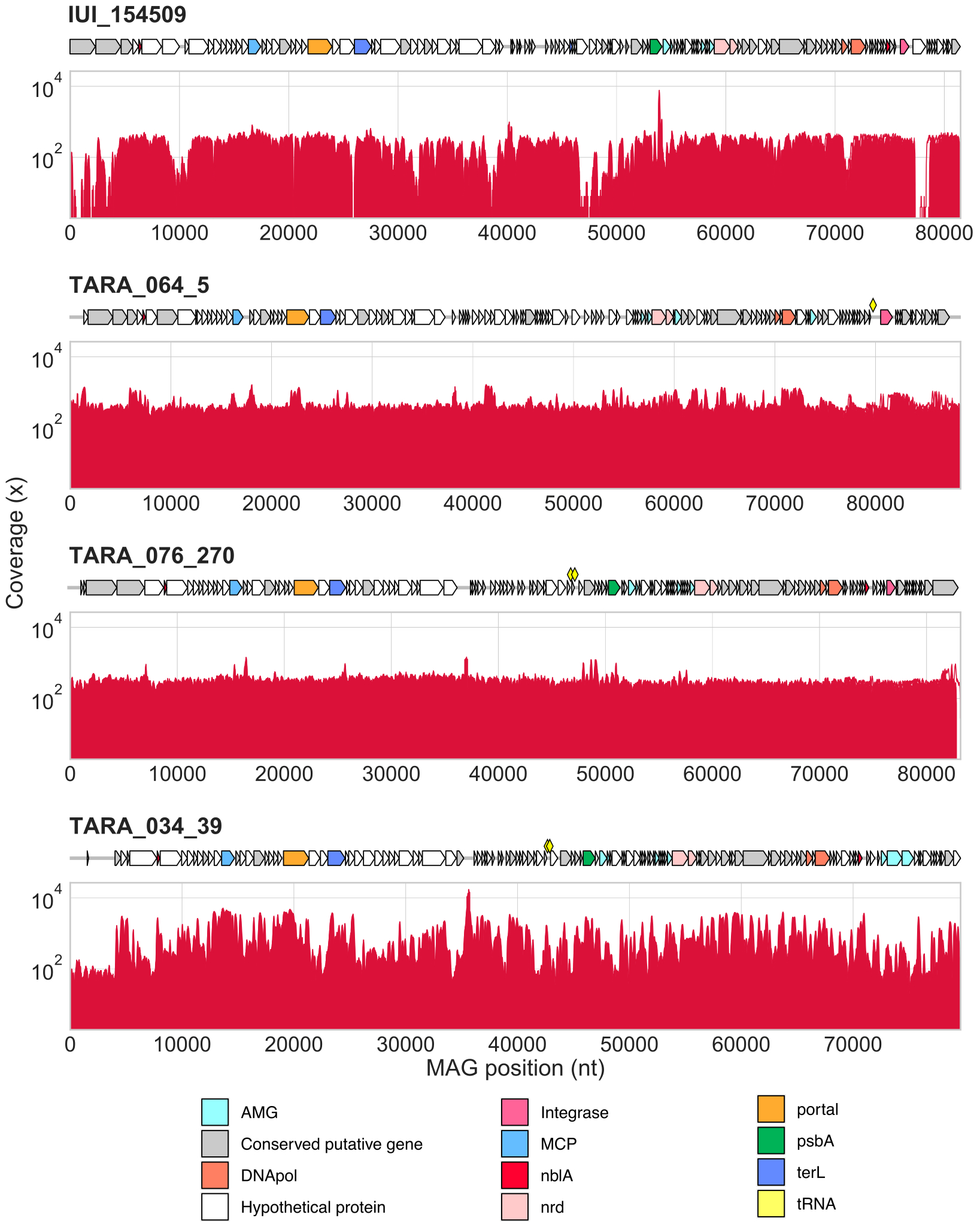

### Supplemental Figure 4

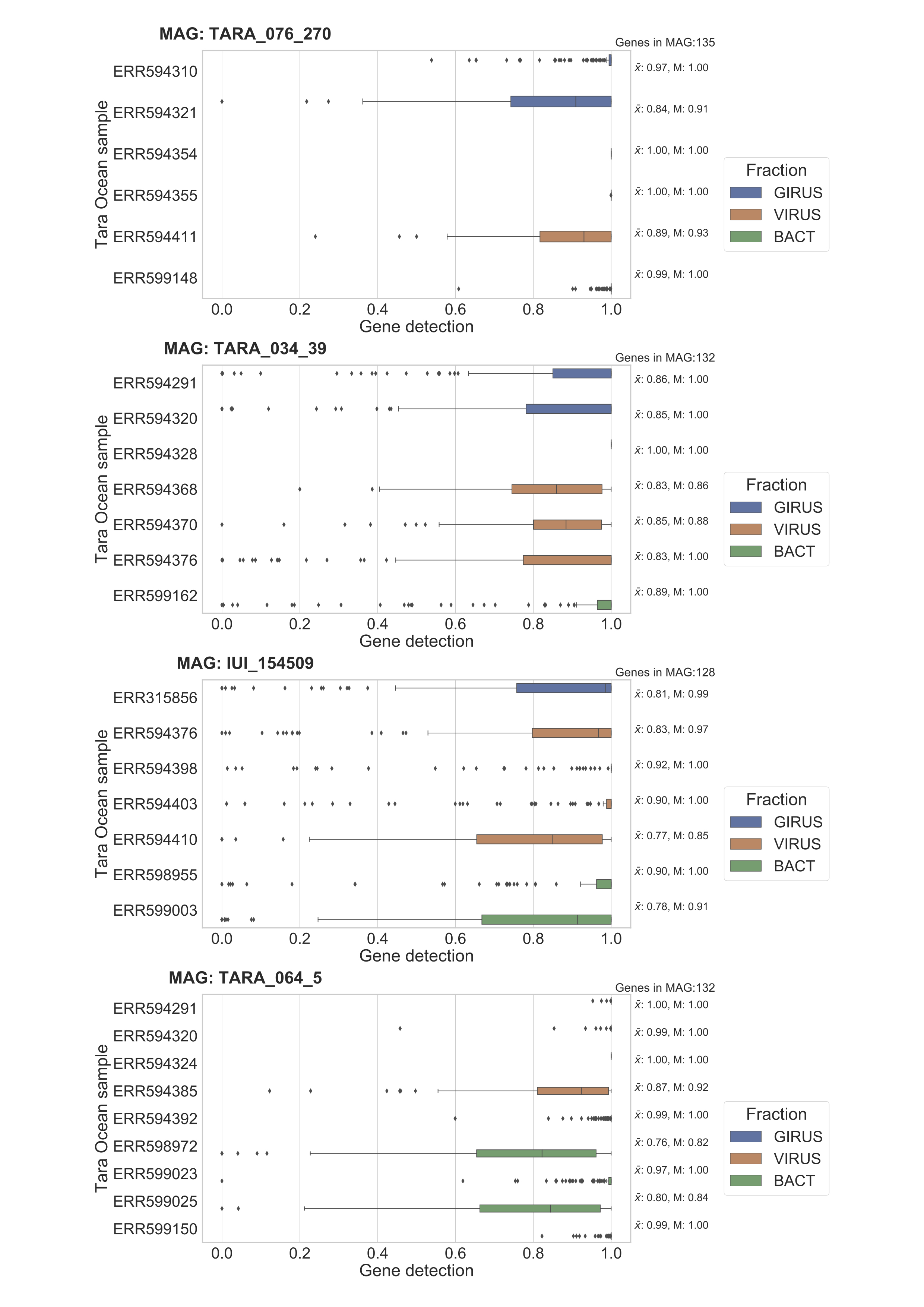
